# Technology-independent estimation of cell type composition using differentially methylated regions

**DOI:** 10.1101/213769

**Authors:** Stephanie C. Hicks, Rafael A. Irizarry

## Abstract

**Background:** High-resolution genome-wide measurement of DNA methylation (DNAm) has become a widely used assay in biomedical research. A major challenge in measuring DNAm is variability introduced from intra-sample cellular heterogeneity, which is a convolution of DNAm profiles across cell types. When this source of variability is confounded with an outcome of interest, if unaccounted for, false positives ensue. This is particularly problematic in epigenome-wide association studies for human disease performed on whole blood, a heterogeneous tissue. To account for this source of variability, a first step is to determine the actual cell proportions of each sample. Currently, the most effective approach is based on fitting a linear model in which one assumes the DNAm profiles of the representative cell types are known. However, we can only make this assumption when a dedicated experiment is performed to provide a plug-in estimate for these profiles. Although this method works well in practice, technology-specific biases lead to platform-dependent plug-in profiles. As a result, to apply the current methods across technologies we are required to repeat these costly experiments for each platform.

**Results:** Here, we present a method that accurately estimates cell proportions agnostic to platform by first using experimental data to identify regions in which each cell type is clearly methylated or unmethlyated and model these as latent states. While the continuous measurements used in the linear model approaches are affected by platform-specific biases, the latent states are biologically driven and therefore technology independent, implying that experimental data only needs to be collected once. We demonstrate that our method accurately estimates the cell composition from whole blood samples and is applicable across multiple platforms, including microarray and sequencing platforms.

## 1 Background

DNA methylation (DNAm) is a type of chemical modification occurring at CpG dinucleotide sites that is involved in controlling gene expression and has been shown to play an important role in distinguishing cell lineages [1]. High-throughput DNAm assays have been widely applied among researchers as well as large consortia to further our understanding of basic biology and health implications [2]. However, a major challenge in extracting information from these DNAm datasets is variability introduced from intra-sample cellular heterogeneity observed in samples of heterogeneous cell composition. Specifically, individual cell types encode unique cell type-specific DNAm signatures to distinguish between the cell lineages. Therefore, when we measure DNAm on samples with a heterogeneous cell composition, we actually observe a convolution of the DNAm profiles of each cell type [3]. It is common for variability in cell type proportions to explain most of the observed sample-to-sample variability.

Cell composition induced variability is particularly problematic in epigenome-wide association studies (EWAS) [4] because, due to convenience, these are most frequently performed on whole blood, a highly heterogeneous tissue. In a seminal paper, Houseman et al. (2012) [3] describe a statistical method that accurately estimates the relative proportions of cell type components in whole blood. Jaffe et al. (2014) [5] used this approach to demonstrate that reported age-related changes of blood DNAm profiles [6, 7, 8, 9, 10, 11, 12] could be explained with high levels of confounding between age-related variability and cell composition, demonstrating the importance of accounting for this source of variability.

As the consequential effect of this source of variability started to be recognized, interest in statistical methods for estimating and accounting for intra-sample cellular heterogeneity grew accordingly. There are currently two major types of approaches. The first, originally developed by Houseman et al. (2012) [3], assumes that the observed heterogeneous blood profiles are a linear combination of the cell type-specific DNAm profiles, assumes these DNAm profiles are known, and then estimates the unknown proportions using a standard estimation procedures. To be able to assume cell type-specific DNAm profiles are known, a rather complex experiment, in which cells of the same cell type are sorted and then used to obtain high-throughput measurements of the reference samples, is conducted. Methods that make use of the sorted cell type-specific DNAm profiles are referred to as *reference-based*. Alternatively, other methods that do not use external reference profiles, referred to as *reference-free* methods, have been developed for DNAm data [13, 14] and for more general types of data such as Surrogate Variable Analysis (SVA) [15], or Remove Unwanted Variability (RUV) [16] to account for batch effects [17].

Reference-based approaches have been shown to greatly outperform reference-free procedures [18]. Here, we consider them to be the state of the art. However, in this paper, we demonstrate that a limitation of reference-based approaches is the presence of a technology-specific bias, which can influence the estimates of cell composition; namely, when using cell type-specific DNAm profiles measured using one technology, for example a microarray platform, to estimate the cell type proportions in samples measured from another technology, for example a sequencing platform. Here, we introduce a statistical method, referred to as *methylCC*, that removes this technical bias using a latent variable model, and accurately estimates the the cell composition in a platform-agnostic manner. To achieve this, we identify regions of the genome in which each cell type are either clearly methylated or unmethylated, and we model these as latent states. These latent states are biologically driven and therefore technology-independent, which allows us to estimate binary, platform-independent profiles that can be successfully be applied across technologies. To study the improvements in estimates of cell composition using methylCC, we evaluated the difference between the true and estimated proportions of cell types with a Monte Carlo simulation. Specifically, we studied how using cell type-specific DNAm profiles measured on a microarray platform to estimate the cell type proportions in samples measured on a sequencing platform can lead to inaccurate estimates of cell composition. Furthermore, we demonstrate how our platform-agnostic approach provides an overall improvement in estimates of cell composition. Although due to the availability of data all our examples are from whole blood, the approach can be generalized to other tissues.

## 2 Results

Consider a set of high-throughput data *Y*_*ij*_ representing a heterogeneous tissue sample, such as whole blood, from *i*∈(1, …, *N*) individuals containing DNAm measurements at CpG sites *j*(1, …, *J*). Suppose the heterogeneous tissue is a combination of *K* cell types, which we index with *k*(1… *K*). Houseman et al. (2012) [3] proposed the following statistical model to estimate the proportions of *K* cell types in whole blood DNAm samples, for each individual *i*:

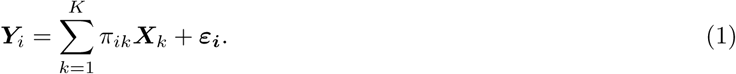

Here *π*_*ik*_ represents the proportion of cell type *k* in individual *i*, which is the parameter of interest. The ***X***_*k*_ represents the *k*^*th*^ cell type-specific DNAm profile with measurements at the same *J* CpGs sites as ***Y***_*i*_. The measurement error and other unexplained biological variability is represented by ***ε***_*i*_. The cell type proportions for individual *i* are assumed to be nonnegative, *π*_*ik*_ ≥ 0, and sum to 1, 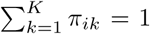. To develop a practical tool, Houseman et al. (2012) [3] sorted whole blood samples into *K* = 6 cell types that make up the majority of this tissue and obtained a DNAm profile for each cell type. They used Illumina’s HumanMethylation27 BeadChip (Illumina 27K), which measures DNAm at approximately 27,000 CpG sites [19]. This experimental data provided plug-in estimates for the cell type-specific DNAm profile, ***X***_*k*_, and with these in place then they estimated the *π*_*ik*_ using a constrained least square algorithm. Soon after the development of this method, Illumina released a new platform that measured approximately 450,000 CpG sites: the HumanMethylation450 BeadChip (Illumina 450K) [20]. Jaffe et al. (2014) [5] leveraged publicly available data of sorted cell types measured with this new Illumina 450K platform [1] to implement the Houseman et al. (2012) method [3].

Although the Illumina 450K microarray platform has been the most widely used platform, two new sequencing technologies are being increasingly adapted by the research community: Whole-genome Bisulfite Sequencing (WGBS) and Reduced Representation Bisulfite Sequencing (RRBS) [21]. Furthermore, Illumina has recently released a new version of their BeadChip, which measures approximately 850,000 CpG sites. However, similar experiments with sorted cells are not yet available from these new platforms, which implies we do not have plug-in estimates for ***X***_*k*_ on these platform technologies. Currently, the only way the Houseman et al. (2012) approach [3] can be applied to DNAm data measured on these new platforms is by assuming that the cell type-specific DNAm profiles ***X***_*k*_ derived for the 450K platform applies to others. Here, we show this assumption does not hold.

### 2.1 Across platforms estimates are inaccurate

To determine if the Houseman et al. (2012) method [3], as implemented by Jaffe et al. (2014) [5], which was specifically developed for the Illumina 450K array platform, is applicable across platforms, we obtained a dataset for which each of 12 whole blood samples were run on both the Illumina 450K and RRBS platforms (referred to below as the *two-platform dataset*). We applied the Houseman method to the whole blood samples in the *two-platform dataset* and expected similar cell composition estimates for each individual across platforms as these were the same whole blood samples just measured on two platforms. Because the Houseman method has been shown to provide reliable cell composition estimates for DNAm data measured on the Illumina 450K platform, in this specific case we considered the cell composition estimates from the Houseman method to be the gold-standard or ground truth, as done by Rahmani et al. (2017) [22]. However, we found that the resulting cell composition estimates between the *N* = 12 whole blood samples measured on the Illumina 450K and RRBS platforms did not agree (Figure 1).

**Figure 1:**
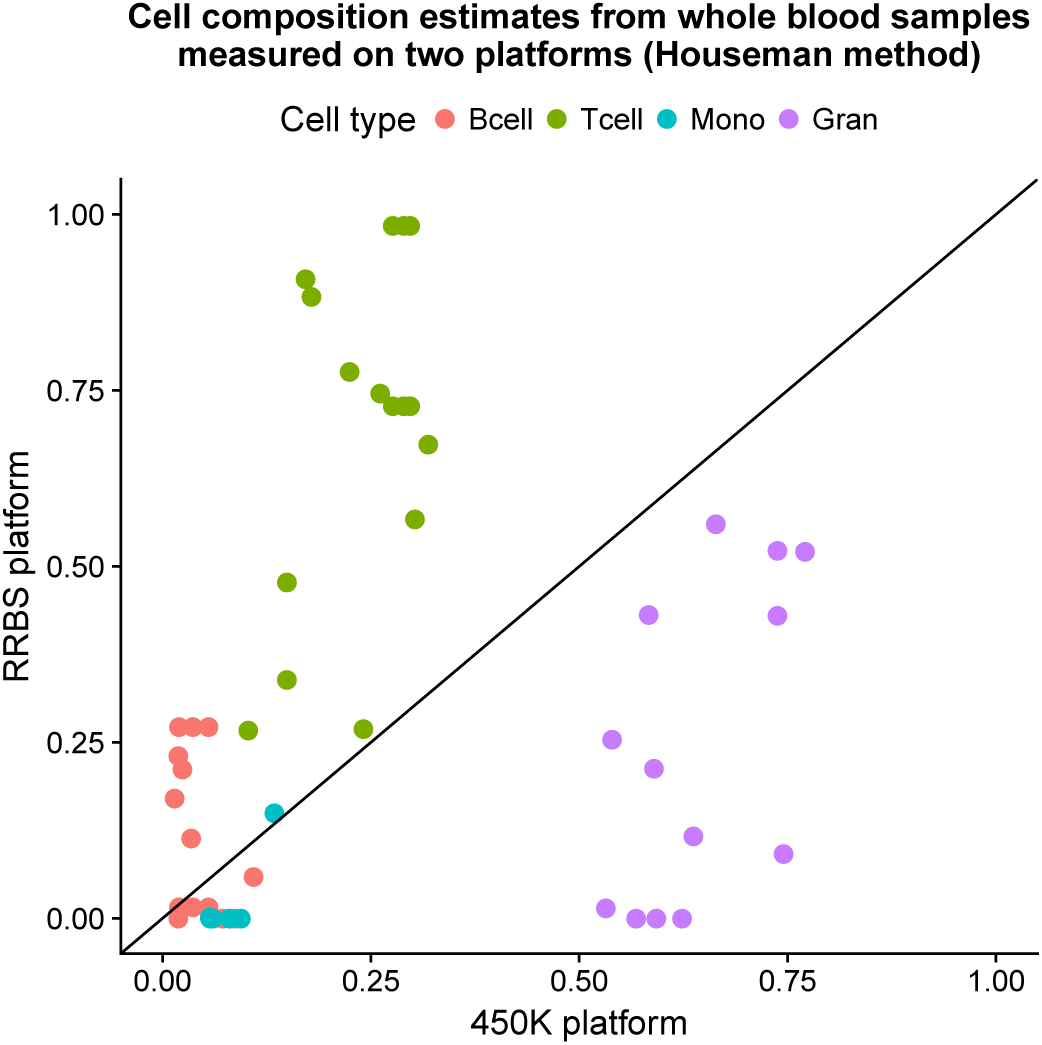
Across platforms estimates are inaccurate. Cell composition estimates (*K* = 4 cell types) from *N* = 12 whole blood samples (*two-platform dataset*) measured on the Illumina 450k microarray platform (x-axis) and the RRBS platform (y-axis). The statistical method proposed by Houseman et al. (2012) [3] was used to estimate the cell composition.

To determine the cause of this disagreement, we examined this dataset more closely and found two limitations with the Houseman approach when applied to technologies other than the Illumina microarrays: (1) DNAm measurements vary across platforms and (2) different platforms measure different CpGs. These two limitations are discussed in the following two sections, respectively. We then describe a statistical solution to overcome these two limitations using a general latent class model to estimate the cell composition of heterogeneous samples agnostic to platform technology. We also provide a software implementation of our method available at https://github.com/stephaniehicks/methylCC.

### 2.2 DNAm measurements vary across platforms

The first limitation is that there is a platform-dependent bias. We can observe this bias by simply plotting and comparing the raw DNAm measurements using the 12 whole blood samples in the *two-platform dataset*. We commonly observe genomic regions in which both platforms seem to indicate a change from unmethylated to methylated states, but the observed DNAm levels differ substantially across platforms (for example, Figure 2A). A more systematic demonstration is obtained by first using the reference cell sorted dataset [1] to identify regions that are clearly unmethylated in all purified cell types and regions that are clearly methylated in all purified cell types, and then plotting the empirical DNAm distribution across all whole blood samples within these regions for both platforms (Figure 2B) and noting the different distributions. We note in particular that observed DNAm levels measured on RRBS tends to have values closer to 0 and 1, compared to the Illumina 450K array attenuating these values away from the edges.

**Figure 2:**
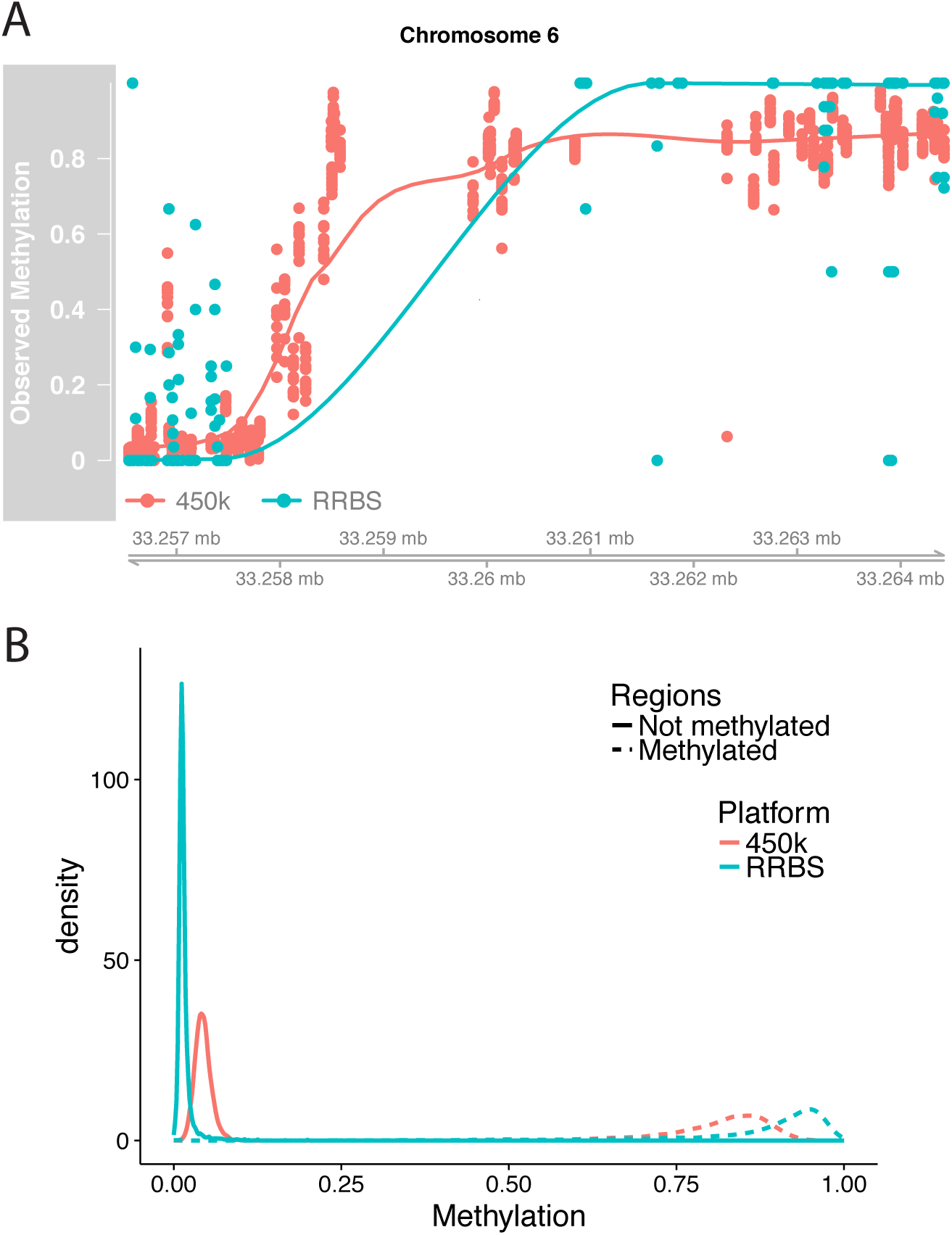
Evidence for platform-dependent bias. (A) Observed DNAm levels from one region in same *N* = 12 individuals (GEO Accession GSE95163) measured on two platform technologies: Illumina 450k (red) and RRBS (blue). (B) Density of DNAm levels measured on the 450k platform (red) and RRBS platform (blue) in regions that are either methylated (dashed line) or not methylated (solid line). Regions were identified by searching for regions in purified whole blood cell types from Reinius et al. (2012) [1] that appeared either methylated or not methylated in all six purified cell types.

### 2.3 Different platforms measure different CpGs

The second limitation is that different platforms measure different CpGs. The human genome contains over 20,000,000 CpGs sites and each platform includes a subset of these which, for logistical reasons, differs across platforms. For example, RRBS [21] uses restriction enzymes to enrich for the areas of the genome that have a high CpG content, while the Illumina 450k platform can build probes for any CpG and selects CpG sites that are more uniformly distributed across the genome. Therefore, to apply the Houseman model to samples measured on platforms other than the Illumina 450K array, we have to restrict ourselves to the intersection of the CpGs measured Illumina 450K array and the alternative platform, because the *j*^*th*^ CpG in the whole blood sample ***Y***_*i*_ must match the *j*^*th*^ CpG in the cell type-specific DNAm profile ***X***_*k*_. As a result, in our 12 RRBS samples we only have measurements from 91 of the 600 CpGs in the cell type-specific DNAm profiles used by the 450K implementation of the Houseman method. This results in a loss of power since informative cell type-specific CpGs may be left out (for example Figure 3).

**Figure 3:**
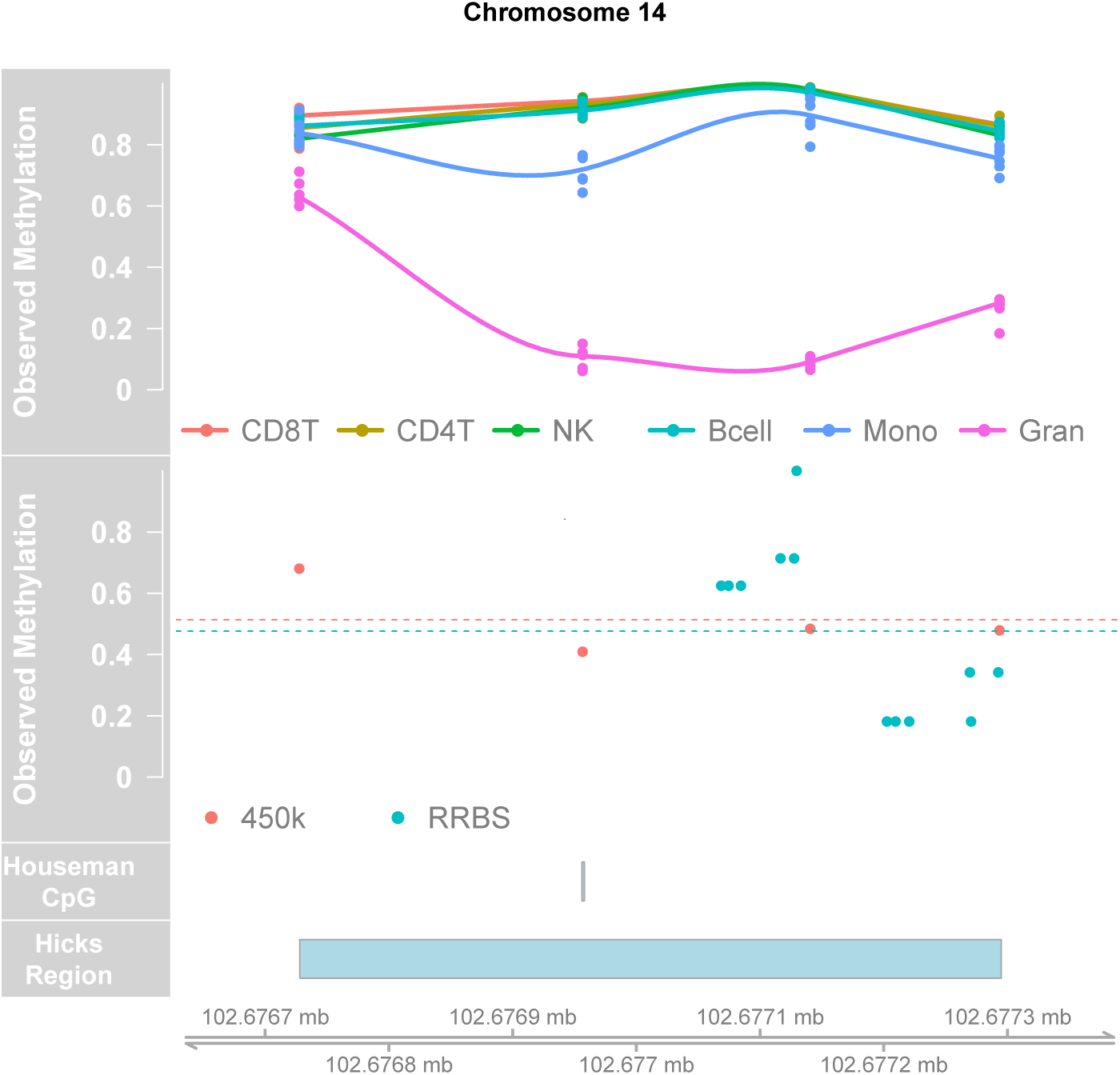
Example of a cell type-specific CpG measured on the 450k microarray platform, but not measured on the RRBS platform. The cell type-specific CpG (Houseman CpG) and the cell type-specific region of CpGs (Hicks Region) were both identified using the purified CD8T, CD4T, Natural Killer (NK), Bcell, Monocytes (Mono) and Granulocytes (Gran) cell types from [1] (top panel). In the whole blood DNAm data from one individual (GEO Accession GSE95163) measured on two platforms (450k and RRBS), the cell type-specific CpG from Houseman et al. (2012) [3] (Houseman CpG) is not measured on the RRBS platform (second panel). However, using a cell type-specific region (Hicks Region), we are able to measure the methylation level averaged across the region in both the 450k and RRBS platform for this sample (dotted lines).

### 2.4 methylcc: estimate cell composition in DNAm samples agnostic to platform technology

To adjust for the platform-specific biases, we introduce a model that accounts for these biases directly and models methylation states using latent variables. To account for the fact that different platforms measure different CpG sites, we model the latent classes at the region level rather than the CpG level. Specifically, we propose the following statistical model:

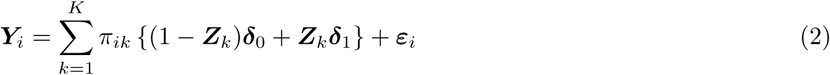

where ***Y***_*i*_ is the observed DNAm level in the heterogeneous tissue, in this case whole blood, for the *i*^*th*^ individual *i* ∈(1, …, *N*), but now measured DNAm levels in *R* genomic regions *r* ∈(1, …, *R*), as opposed to *J* individual CpGs in the Houseman model. Similar to the Houseman model, *π*_*ik*_ represents the proportion of cell type *k* in individual *i*, which is the parameter of interest. In addition, we assume that the cell type proportions for individual *i* are nonnegative, *π*_*ik*_ ≥ 0, and sum to 1, 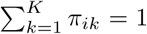. Here, ***Z***_*k*_ = (*Z*_1*k*_, …, *Z*_*Rk*_) is a vector of latent variables for the *k*^*th*^ cell type where each latent variable, *Z*_*rk*_, is an indicator that is equal to 1 if the region *r* is methylated in cell type *k* and 0 otherwise. The platform-specific biases are represented with random effects ***δ***_0_ = (*δ*_0,1_, …, *δ*_0,*R*_) and ***δ***_1_ = (*δ*_1,1_, …, *δ*_1,*R*_), which are assumed to follow multivariate normal distributions 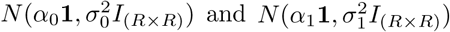, respectively. Measurement error and other unexplained biological variability is represented with ***ε***_*i*_, which we assume follows a multivariate normal distribution *N*(0, *τ*^2^*I*_(*R×R*)_). Note that in our model the random effects ***δ***_0_ and ***δ***_1_ are assumed to be platform-dependent: they represent the technology-dependent bias with different mean and variances in different platforms (Figure 2B). However, the ***Z***_*k*_s are not platform-dependent: they are latent classes determined by biology.

The statistical model in Equation 2 can be thought of as a generalization of Equation 1 if we restrict the Houseman approach to only include CpGs that are either methylated or unmethylated in each cell type. In this case, region *r* would simply be a single CpG site and the *k*^*th*^ cell type-specific DNAm profile, ***X***_*k*_, would be defined by *X*_*rk*_ = *δ*_0,*r*_ if region *r* is unmethylated and *X*_*rk*_ = *δ*_1,*r*_ if region *r* is methylated.

A significant advantage of our model is that instead of directly measuring the cell type-specific DNAm profiles, ***X***_*k*_, for each platform, we account for region-to-region variability using a latent random variable and therefore do not need to measure it directly with each new platform. Instead, all we need is to identify *R* regions for which each *k*^*th*^ cell type is either clearly methylated (*Z*_*rk*_ = 1), or not methylated (*Z*_*rk*_ = 0) for *r* ∈ (1, …, *R*). We define ***Z*** to be the matrix with entries *Z*_*rk*_ in the *r*-th row and *k*-th column, which needs to be full rank for the parameters of interest, *π*_*ik*_ to be identifiable. Because ***Z*** is entirely determined by biology, not by the platform technology, we only have to identify these regions once for each type of heterogeneous (biological) sample. This requires experimental data from cell sorted samples measured on only one platform. To demonstrate the utility this approach for estimating cell composition in whole blood samples, we searched for these genomic regions in the purified cell type data described in [1], which were measured on the Illumina 450K array platform. This dataset includes Bcells, monocytes, granulocytes, CD8T, CD4T and natural killer (NK) cells. We combined the last three into ‘Tcells’ because the DNAm profiles were too similar and it was not possible to form a full rank matrix if included. We identified *R* = 212 regions satisfying our criteria (Figure 4). Finally, with the *R* regions in place, the estimation of the proportion of cell types, *π*_*ik*_, reduces to a missing data problem. We use an EM algorithm with a constrained linear model to estimate the parameters 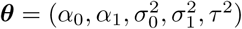 and ***π**_i_* = (*π*_*i*1_, …, *π*_*iK*_) for individuals *i* ∈ (1, …, *N*) (see the Materials and Methods Section for complete details on estimation procedure).

**Figure 4:**
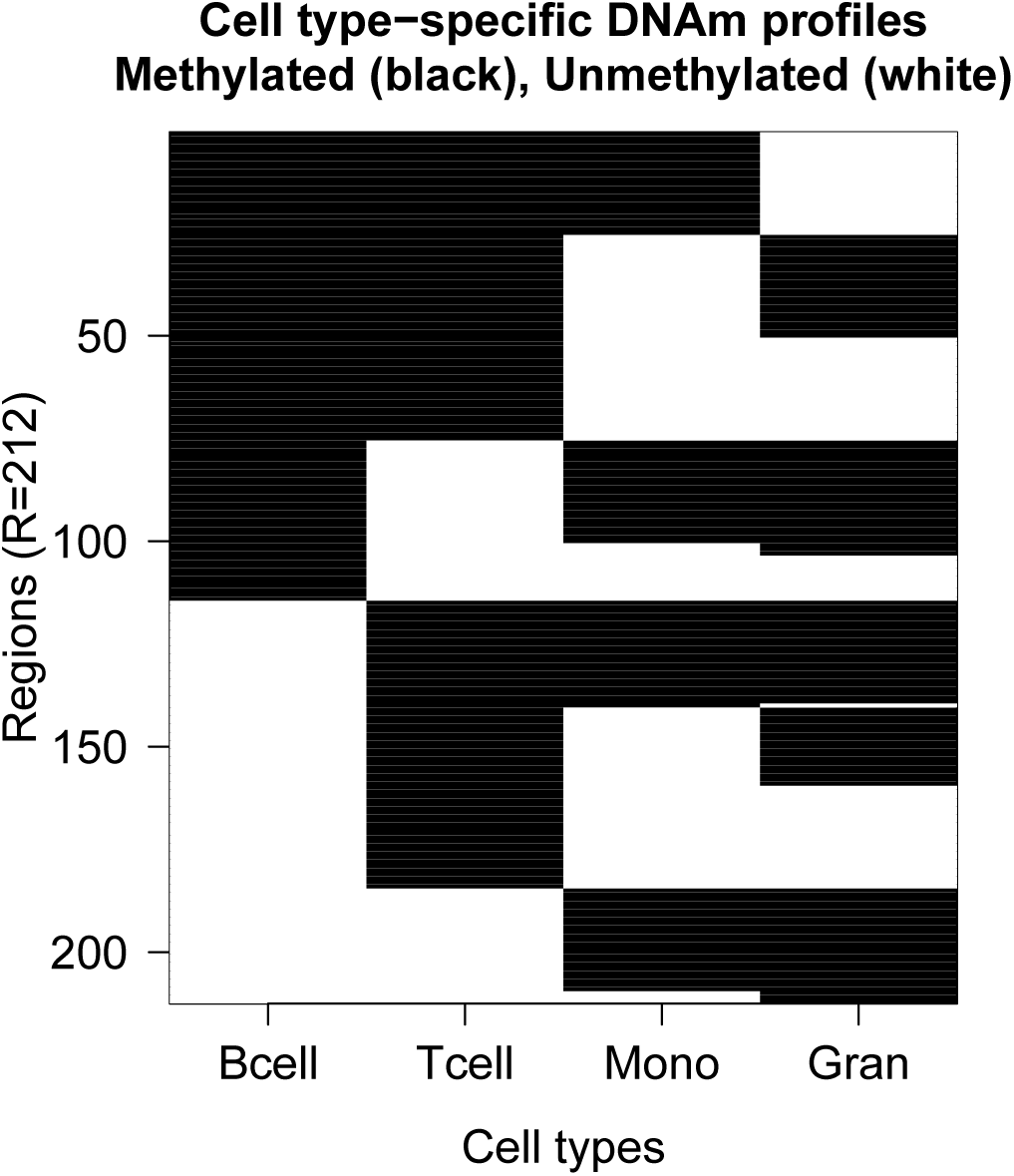
Cell type-specific DNAm profiles in whole blood. The image contains the *R* = 212 informative genomic regions containing regions that are either methylated (black) or unmethylated (white) in Bcells, Tcells, Monocytes (Mono) and Granulocytes (Gran).

### 2.5 methylcc improves estimates of cell composition of DNAm samples measured on other platform technologies

To demonstrate the improvements in the estimates of cell composition provided by our platform-agnostic approach, we applied our method to the *two-platforms dataset*. Specifically, we fit our model to the 12 whole blood samples measured on both the Illumina 450K array and RRBS platforms. Similar to Figure 1, we considered the estimates of cell composition from the Houseman model in the Illumina 450K samples to be the gold-standard reference. In Figure 1, we demonstrated that directly applying the Houseman approach [3], as implemented by Jaffe et al. (2014) [5], to the RRBS data led to biased cell composition estimates. However, our new approach substantially improves estimates of cell composition (Figure 5).

**Figure 5:**
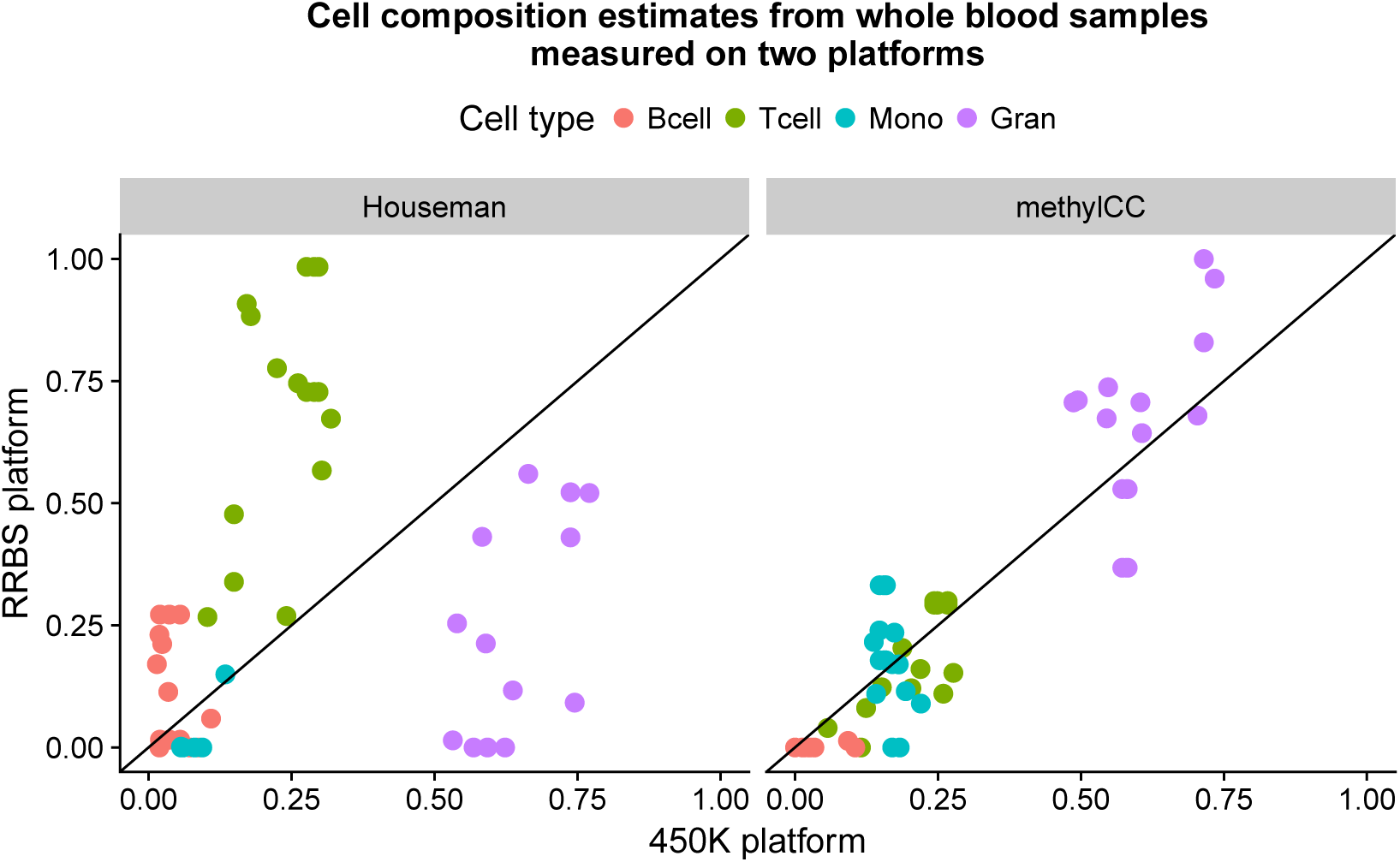
methylCC: A latent variable model with region-specific and platform-dependent random effects improves estimates of cell composition. Cell composition estimates from *N* =12 whole blood samples (*K*=4 cell types) measured on the Illumina 450k microarray platform (x-axis) and the RRBS platform (y-axis). Two methods were used to estimate the cell composition: (1) the model proposed by Houseman et al. (2012) [3] that was developed for samples measured on the Illumina 450k microarray platform (left), and (2) our proposed method that is independent of platform technology (right).

Furthermore, we evaluate the performance of our platform-agnostic approach with the goal of estimating the proportion of cell types in heterogeneous tissue samples. Here, we performed a Monte Carlo simulation study to illustrate the improvements in estimates of cell composition by our platform-agnostic approach compared to the Houseman approach for heterogeneous samples measured on a sequencing platform (described in detail in Materials and Methods Section). For the simulations study, we created cell type-specific DNAm profiles for a microarray platform, 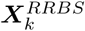, and a sequencing platform, 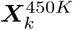, by simulating platform-dependent random effects with different means and variances (Figure S1A). Then, we simulate whole blood samples with a relative proportion of cell types ***π***_*i*_ and measurement error ***ε***_*i*_ to create the observed DNAm level in whole blood samples measured on in the 450k array platform 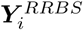 and the RRBS platform 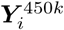. We estimate the cell composition in the whole blood samples measured on both platform using the reference-based Houseman method and our platform-agnostic method. Then, we evaluate the difference between the true and estimated proportion of cell types using either our approach or the Houseman approach.

**Figure S1:**
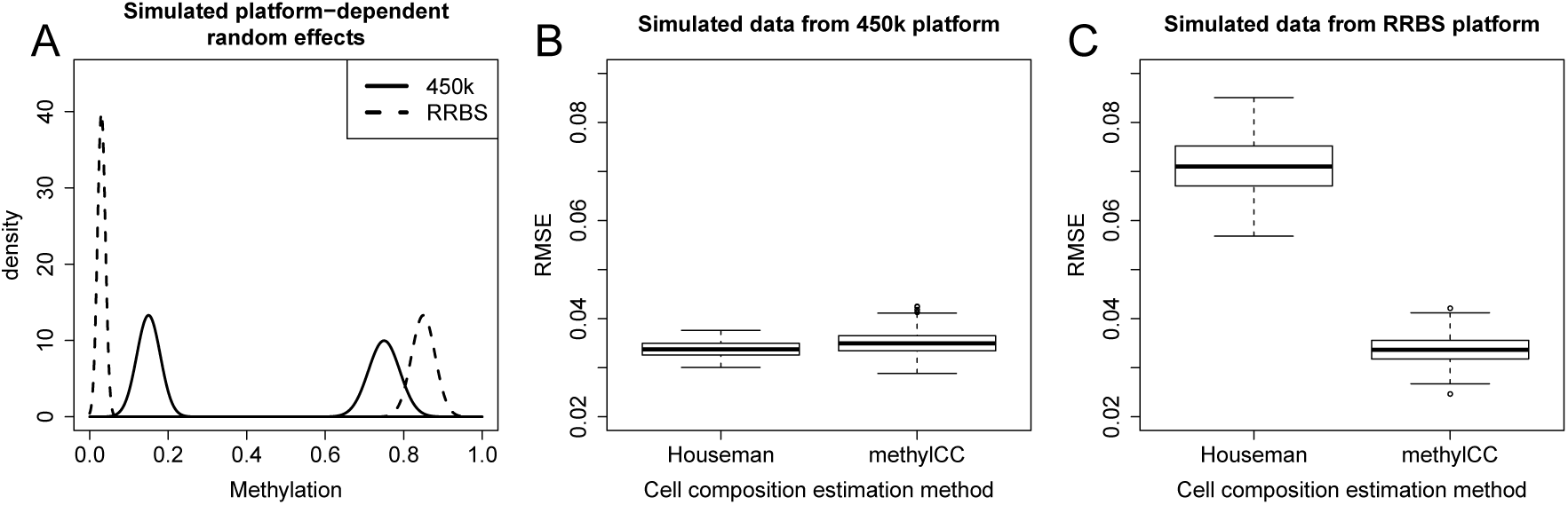
Simulation study illustrating improvement in RMSE by accounting for platform-dependent effects. (A) Simulated platform-dependent random effects, similar to the platform-dependent effects observed in Illumina 450K and RRBS data (Figure 2B). Root mean squared error (RMSE) averaged across cell types for whole blood DNAm samples measured on (B) the Illumina 450k array platform and on (C) the RRBS platform. Within each plot, the RMSE is shown for the reference-based method from Houseman et al. (2012) [3] and our proposed model (methylCC).

For whole blood samples measured on the 450K array platform, we found the Houseman approach, which was specifically developed for the array platform, and our approach perform similarly (Figure S1B). However, for whole blood samples measured on a sequencing platform, our platform-agnostic model results in significantly improved estimates of cell composition (Figure S1C). This is because our model accounts for the platform-specific biases directly and models methylation states using latent variables.

### 2.6 methylcc accurately estimates cell composition of DNAm samples measured on Illumina 450K array

To validate the results of our simulation study using whole blood samples measured on the Illumina 450K array platform, we compared the cell composition estimates from our model to the cell composition estimates from the Houseman model using publicly available DNAm whole blood samples measured on the Illumina 450K array platform. Similar to before, we considered the cell composition estimates from the Houseman model to be the “gold standard” for the purposes of this assessment because the Houseman model was specifically designed for the Illumina 450K array platform and it has been previously considered as a “gold standard” [22]. We found that our platform-agnostic approach closely matches the reference-based approach using *N* = 689 whole blood samples from Liu et al. (2013) [23] (Supplemental Figure S2).

## 3 Discussion and Conclusions

We have developed a latent variable model with region-specific and platform-dependent random effects to accurately estimate the cell composition in DNAm whole blood samples measured from any platform technology. By using informative genomic regions that are either methylated or unmethylated for each purified cell type, we illustrated how we can estimate the cell composition across platform technologies as cell types preserve their methylation state in regions independent of platform, despite observed measurements being platform-dependent. Note that, our current model assumes that the random effects and measurement error are normally distributed. Although these assumptions were a practical approximation that led to an improvement for RRBS data, with WGBS data the model may need to be generalized to other distributions since data sets produced with this platform may produce count data for which negative binomial models may be more appropriate.

We have demonstrated the utility of our method by applying it to real and simulated whole blood samples measured on a sequencing platform. We illustrated how our method significantly improves the estimates of the cell composition in whole blood samples measured on a sequencing platform compared to the reference-based method, because our model accounts for the platform-specific biases directly and models methylation states using latent variables. Given that sequencing platform technologies are poised to become more widely used for studies measuring DNAm in whole blood, this suggests that our method is an needed contribution.

**Figure S2:**
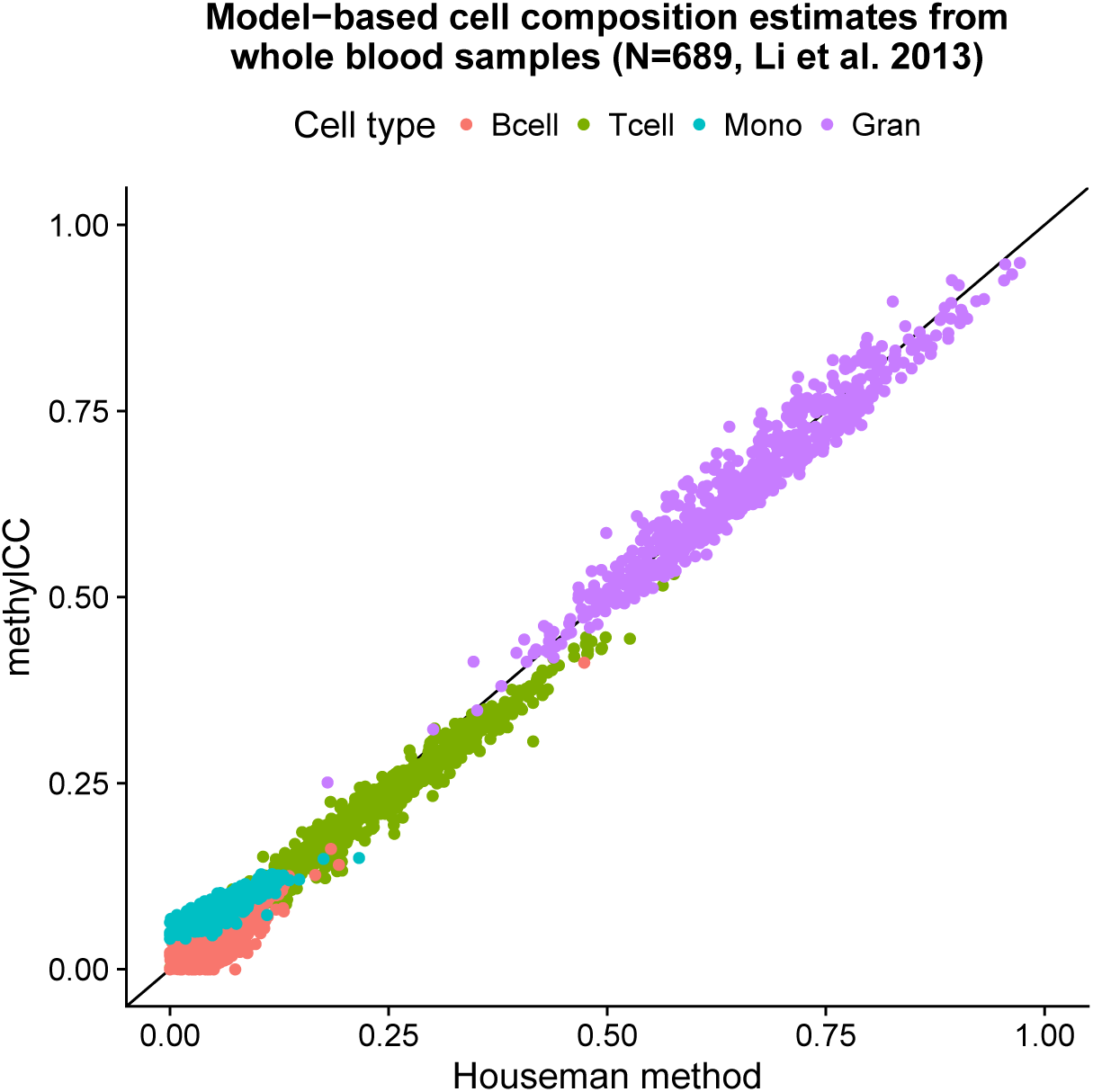
Platform-agnostic approach closely matches the reference-based approach using data measured on the Illumina 450K platform. Cell composition estimates (*K*=4 cell types) for *N*=689 whole blood samples from Liu et al. (2013) [23] measured on the Illumina 450K array platform. Here, the cell composition estimates from the Houseman method (x-axis) are considered the “gold standard” because this method was specifically designed for the Illumina 450K array platform. We compared this reference-based approach to cell composition estimates from our platform-agnostic approach (methylCC) (y-axis).

## 4 Materials and Methods

### Using cell sorted experimental data to identify informative genomic regions in *Z*

Cell sorted experimental data is needed to identify *R* informative genomic regions for which the *k*^*th*^ cell type is either clearly methylated (*Z*_*rk*_ = 1) or not methylated (*Z*_*rk*_ = 0) for regions *r* ∈(1, …, *R*). This is step is done only once for each type of heterogeneous (biological) sample such as whole blood and does not depend on the platform technology. In addition, this matrix ***Z*** needs to be full rank for the parameters of interest, *π*_*ik*_ to be identifiable.

In application for estimating the cell composition in whole blood samples, we used cell sorted data described in [1], which were measured on the Illumina 450K array platform. This dataset includes six biological replicates for each of the six purified cell type (Bcells, monocytes, granulocytes, CD8T, CD4T and natural killer (NK) cells). We used the `bumphunter` Bioconductor package [24] to identify differentially methylated regions (DMRs) across cell types. For example, to search for DMRs such that the six granulocytes samples are unmethylated and the other cell types are methylated (Figure 3), we fit a linear model *Y*_*ij*_ = *β*_0_(*l*_*j*_) + *β*_1_(*l*_*j*_)*X*_*j*_ + *ε*_*ij*_ at each *j*^*th*^ genomic position (or CpG site) where *Y*_*ij*_ represents observed DNAm level in the *i*^*th*^ biological replicate for a purified cell type at position *j* with a covariate of interest, *X*_*j*_, (for example *X*_*j*_ = 0 for granulocytes and *X*_*j*_ = 1 for other cell types). Then, we searched for regions of CpGs such that *β*_1_(*l*)≠ 0. For more details on identifying DMRs, we refer the reader to [24, 25].

We searched for regions that were not overlapping so they would be considered independent observations. In certain pairwise cell type comparisons, the only regions found contained just one CpG, however we prioritized regions with more than one CpG whenever possible. Also, we combined three cell types (CD8T, CD4T and NK cells) into one cell type ‘Tcells’ when searching for DMRs because the DNAm profiles were too similar and it was not possible to form a full rank ***Z*** matrix. Following these steps, we identified *R* = 212 regions satisfying our criteria (Figure 4).

In the next section, we describe our estimation procedure to obtain the cell composition estimates, ***π***_*i*_ = (*π*_*i*1_, …, *π*_*iK*_), and we note that we assume these regions ***Z*** are known here. This is because if we fit the model only to these regions, then the estimation procedure reduces to a missing data problem with random effects ***δ***_0_ and ***δ***_1_.

### Estimation procedure

Using the *R* = 212 informative genomic regions identified above, we estimate the parameters of interest, namely the proportion of cell types ***π***_*i*_ = (*π*_*i*1_, …, *π*_*iK*_) for the *i*∈(1, …, *N*) individuals, and the parameters 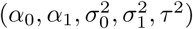 in the proposed latent variable model (Equation 2) using an EM algorithm with constraints 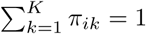 and *π*_*ik*_ ≥ 0 for all *k*.

#### Obtain initial starting values *θ*^(0)^ and 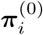 at step *t* = 0

To obtain initial starting values for the 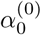 and 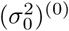 at step *t* = 0, we use the reference cell sorted dataset [1], which has six technical replicates for each cell type, to identify a set of *R*^0^ genomic regions that are clearly unmethylated (*Z*_*rk*_ = 0) in all *K* purified whole blood cell types. In these unmethylated regions, the expected DNAm level is

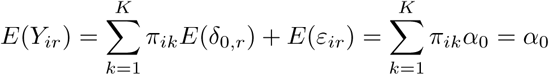

and we use Jensen’s inequality to estimate an upper bound on the variance of *Y*_*ir*_:

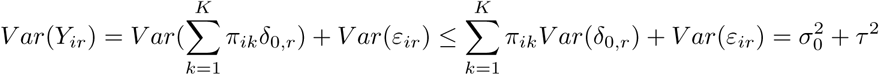

Therefore, we obtain initial starting values

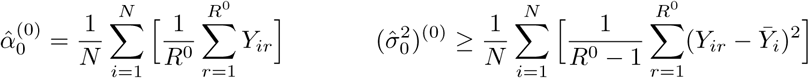

where the measurement error *τ*^2^ is assumed to be small. The argument is similar for the initial starting values of 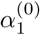 and 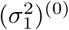 by identifying genomic regions (*R*^1^) where the CpGs are all methylated (*Z*_*rk*_ = 1) for all *K* purified cell types.

To obtain initial starting values for the proportion of cell types, 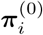 with constraints 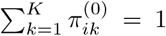 and 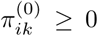 we use the fact that 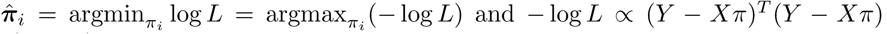. This non-negative least squares (NNLS) problem with constraints is equivalent to the quadratic programming problem 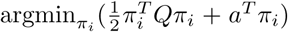 where *Q* = (*X*^*T*^*X*) and *a* = (*−X*^*T*^*Y*) [26]. Therefore, we calculate 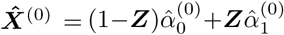 and apply quadratic programming [26] to solve for 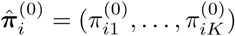. We use the `solve.QP()` function from the R package quadprog [27] to implement the quadratic programming. Finally, to obtain an initial starting value for (*τ*^2^)^(0)^, we calculate

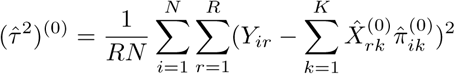

#### EM algorithm to estimate *θ* and *π*

To construct an EM algorithm to obtain maximum likelihood estimates of ***θ*** and ***π***, we define the complete-data vector ***Y***^∗^ = (***Y***, ***δ***_0_, ***δ***_1_) where ***Y*** = (***Y***_1_, …, ***Y***_*N*_) represents the observed DNAm levels for individuals *i* ∈(1, …, *N*) each of length *R* regions. The complete-data likelihood is given by

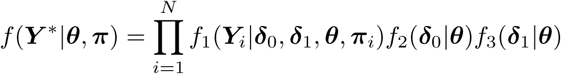

where 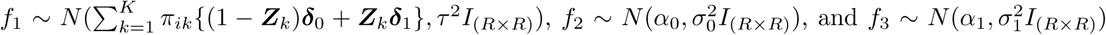. It is easy to show the log of the complete-data likelihood is linear in the following complete-data sufficient statistics: 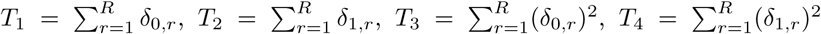 and 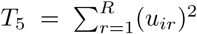 where 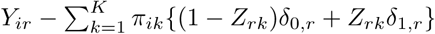.

The EM algorithm alternates between the following two steps:

1. **E-Step**

We can consider the two joint distributions ***Y***^∗^ = (***Y***, ***δ***_0_) and ***Y***^∗^ = (***Y***, ***δ***_1_) separately since ***δ***_0_ and ***δ***_1_ are independent. The joint distributions are also normally distributed

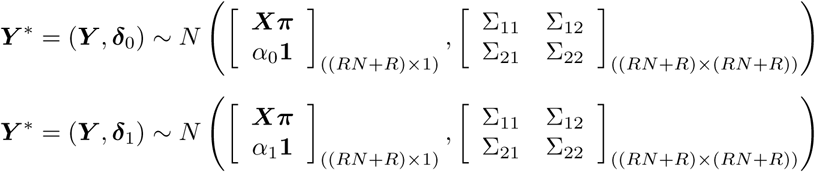

where ***Y*** is a matrix of dimension *R*×*N*, but we convert this into a vector of length *RN*, ***X*** = (1−***Z***)*α*_0_ + ***Z**α*_1_ is an *R* × *K* matrix and ***π*** is a *K* × *N* matrix. We convert the ***Xπ*** matrix into a vector of length *RN*. To derive the conditional distributions of ***δ***_0_|***Y*** and ***δ***_1_|***Y***, we use Theorem 3.2.3 and 3.2.4 in [28]:

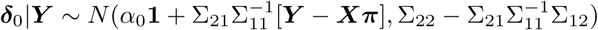

where

- ***X*** = (1 − **Z**)*α*_0_ + ***Z***α_1_ is an *R* × *K* matrix. ***π*** is a *K* × *N* matrix.
- Σ_11_ = *Cov*(***Y***) is an *RN* × *RN* covariance matrix with entries

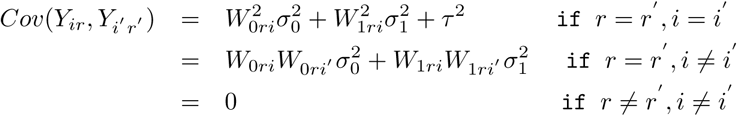

where 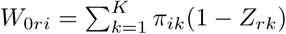 and 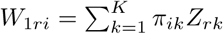
- Σ_12_ = *Cov*(***Y***, ***δ***_0_) is an *RN* × *R* covariance matrix with entries

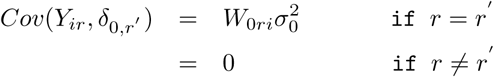 Note: 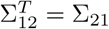
- Σ_22_ = *Cov*(***δ***_0_) is an *R* × *R* matrix with 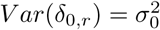 and *Cov*(*δ*_0,*r*_, *δ*_0,*r′*_) = 0

We use the conditional distribution ***δ***_0_|***Y*** to calculate the *t*^*th*^ iteration in the E-Step when computing *E*_*θ*_(*T*_1_|***Y***) and *E*_*θ*_(*T*_3_|***Y***).

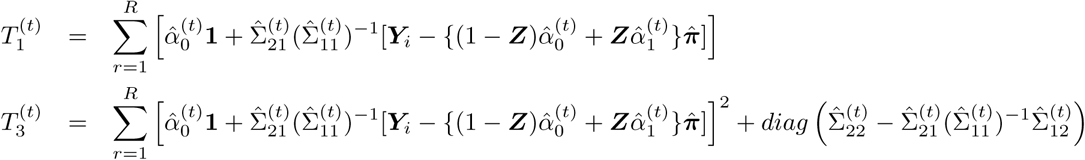

Similarly, we can show

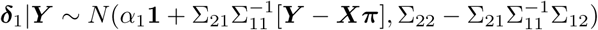

where

- ***X*** = (1 − ***Z***)*α*_0_ + ***Z**α*_1_ is an *R* × *K* matrix. ***π*** is a *K* × *N* matrix.
- Σ_11_ = *Cov*(***Y***) is same as defined above.
- Σ_12_ = *Cov*(***Y***, ***δ***_1_) is an *RN* × *R* covariance matrix with entries

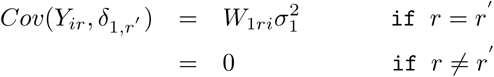

Note: 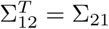
- Σ_22_ = *Cov*(***δ***_1_) is an *R* × *R* matrix with 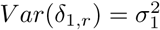 and *Cov*(*δ*_1,*r*_, *δ*_1,*r′*_) = 0

We use the conditional distribution ***δ***_1_|***Y*** to calculate the *t*^*th*^ iteration in the E-Step when computing *E*_*θ*_(*T*_2_|***Y***) and *E*_*θ*_(*T*_4_|***Y***).

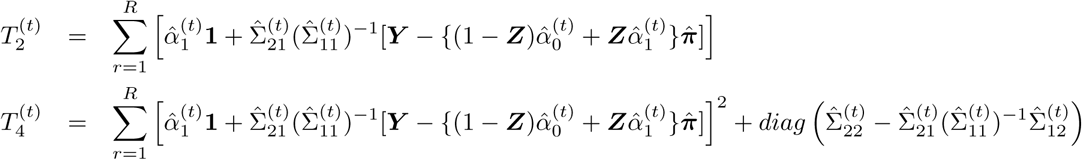

2. **M-Step**

The complete-data maximum likelihood estimates (MLEs) were calculated by using the log of the complete-data likelihood, taking the derivative with respect to the individual parameters, setting the likelihood equal to zero and solving for the MLEs.

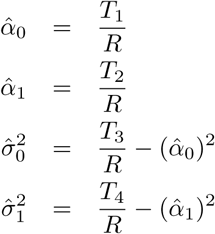

Using these MLEs, we can substitute the sufficient statistics calculated in the E-Step:

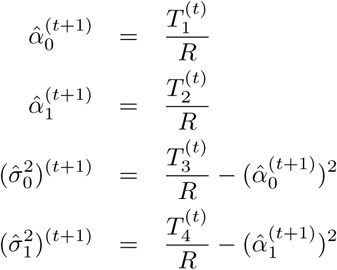

To estimate ***π***_*i*_, we apply quadratic programming [26] (see section on obtaining initial starting values for details) with the constraints 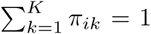 and *π*_*ik*_ ≥ 0 for all *k*. We calculate ***X***^(*t*)^ using the *t*^*th*^ iteration of the conditional expectations *E*_*θ*_(***δ***_0_|***Y***) and *E*_*θ*_(***δ***_1_|***Y***) then apply quadratic programming [26] to solve for 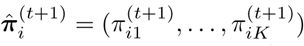. We use the `solve.QP()` function from the R package quadprog [27] to implement the quadratic programming.

Finally, the MLE for *τ*^2^, was calculated by using the log of the complete-data likelihood, taking derivative with respect to *τ*^2^, setting likelihood equal to zero and solving.

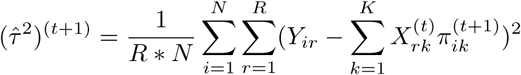

### Publicly available data

We used a publicly available dataset from Liu et al. (2013) [23] measuring the DNAm levels in whole blood samples (*N* = 689) on the Illumina 450K array platform. This dataset from Liu et al. (2013) studied methylation differences between rheumatoid arthritis patients and normal controls (GSE42861). Here, we only considered the normal controls and used the proportion of cell types estimated using the Houseman approach as a gold-standard or the true cell composition in each whole blood sample.

We validated our model by comparing the estimates cell composition of the same whole blood samples measured on two platform technologies from Carmona et al. (2017) [29] (GSE95163): (1) Illumina 450k microarray platform and (2) reduced representation bisulfite sequencing (RRBS) platform. The whole blood samples were derived from ten male individuals resulting in *N*=12 microarray samples and *N* =12 RRBS samples.

### Details for simulation studies

We created platform-dependent cell type-specific DNAm profiles for the *k*^*th*^ cell type 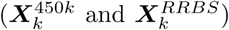 where 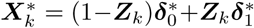 by simulating platform-dependent random effects 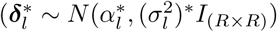 for both *l* = 0,1 (Figure 2B). For each whole blood DNAm sample (*N* = 200), we simulate a relative proportion of cell types (***π***_*i*_) and measurement error (***ε***_*i*_) to create the observed DNAm level in the 450k array platform 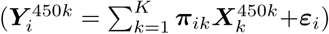 and the RRBS platform 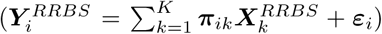. Next, we estimate the cell composition of the 450k array and RRBS samples using both the reference-based Houseman method and our platform-agnostic method. The cell compositions estimates are scaled to sum to 1 if needed. We calculate the cell type-specific *RMSE*_*k*_ as

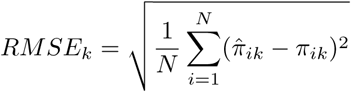

where *π*_*ik*_ is the true cell composition and 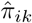 is the estimated cell composition (using either Houseman model or our proposed model) in the *i*^*th*^ sample and *k*^*th*^ cell type. The cell type-specific *RMSE*_*k*_ is averaged across cell types and recorded as the mean RMSE. We repeat the above *n*_*sims*_ = 100 times to calculate the distribution of mean RMSE.

### Software

Software implementing the presented method to estimate the cell composition of whole blood samples measured from DNAm is available as an R package on GitHub (https://github.com/stephaniehicks/methylCC).

## 6 Author’s contributions

SCH and RAI developed the method methylCC. SCH wrote the methylCC R package, analyzed DNAm data and performed the simulation studies. SCH and RAI wrote the manuscript. Both authors read and approved the final manuscript.

